## Supplemental Data for "Interferons and tuft cell numbers are bottlenecks for persistent murine norovirus infection"

Includes:

Figure S1

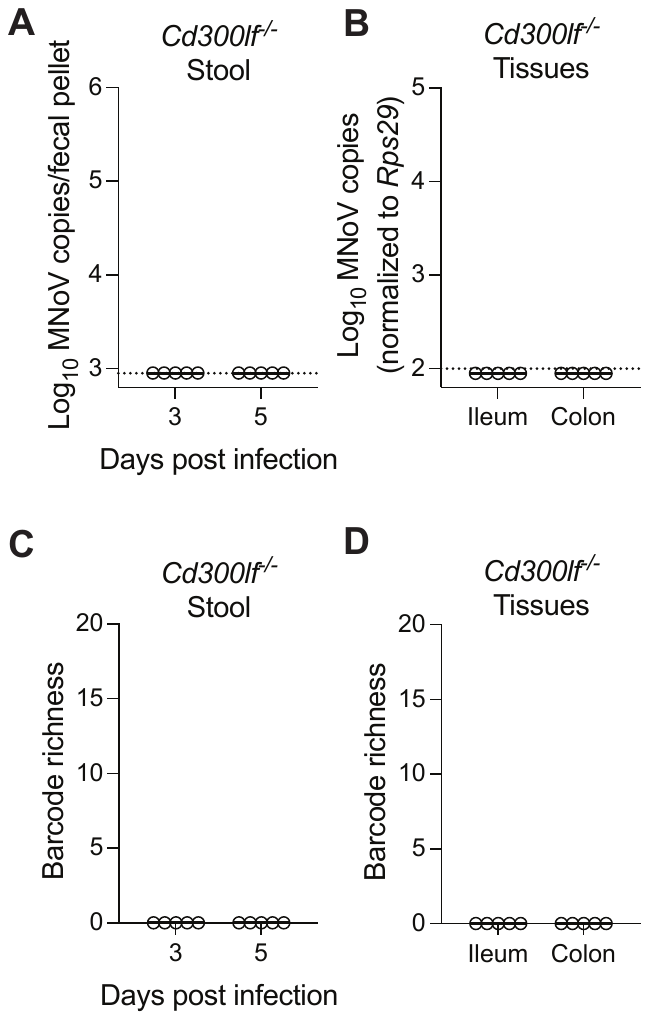

**Supplementary Figure 1: Barcoded CR6 (CR6^BC^) shows similar infectivity to CR6 and maintains barcodes *in vitro* and *in vivo.*** (**A,B**) *Cd300lf^-/-^* mice (N=5) were inoculated with CR6^BC^ and stool viral shedding at 3 and 5dpi (**A**) and tissue viral levels at 5dpi (**B**) were assessed using qPCR. **(C,D)** Barcode richness of *Cd300lf^-/-^* stool (**C**) and tissues (**D**) was determined.
